# Unveiling Eukaryotic Membrane Proteins in High Resolution Using Peptide Solubilization

**DOI:** 10.1101/2025.08.27.672771

**Authors:** Jiahe Zang, Yiting Shi, Weiyu Tao, Xiaoyu Liu, Wenjun Guo, Lei Chen

## Abstract

Integral membrane proteins are vital for numerous biological functions and are typically studied using X-ray crystallography and cryo-electron microscopy (cryo-EM). However, these techniques require the extraction of target membrane proteins from their native membranes using detergents, which might disrupt the lipid environments and alter protein behavior. In this study, we present a novel method for solubilizing membrane proteins using a peptide, thereby eliminating the need for detergents throughout the procedure. We demonstrate that the 4F peptide effectively solubilizes a range of membrane proteins and complexes into peptidiscs, while preserving their functionality and structural integrity. Converting these peptidiscs into nanodiscs further enhances particle homogeneity and facilitates high-resolution structural determination of membrane proteins. Our findings highlight the potential of membrane-solubilizing peptides to advance membrane protein research.

**Highlights:**

- 4F peptide can effectively solubilize eukaryotic membrane proteins
- 4F-peptidisc can be converted to MSP-wrapped nanodisc for high resolution cryo-EM studies
- 4F solubilization can preserve endogenous ligands bound to the target membrane protein

## Introduction

Integral membrane proteins are crucial for various biological processes, including signal transduction, transport, and enzymatic catalysis. Structural studies of these proteins provide essential insights into their mechanisms of action. Typically, the determination of membrane protein structures through techniques such as X-ray crystallography or single-particle cryo-electron microscopy (cryo-EM) necessitates the extraction of the target protein from its native membrane environment using detergents.

Detergents are amphipathic compounds that can be inserted into the lipid bilayer, encapsulating the transmembrane regions of membrane proteins within micelles, which allows for their solubilization in aqueous solutions ^1^. Once solubilized, membrane proteins can be subjected to various purification processes. However, identifying the appropriate detergent for efficient solubilization and stabilization of each specific membrane protein is a challenging task ^2^. Furthermore, detergents can strip away native lipids from membrane proteins, potentially altering the functional behavior of certain proteins ^3^. To address these challenges, researchers have developed methods to remove detergents from solubilized membrane proteins and reconstitute them into structures such as nanodiscs ^4-7^, peptidiscs ^8-10^, or amphipol-wrapped particles ^11^ for biochemical or structural studies. Despite these advancements, the native lipids lost during detergent solubilization may not be fully restored during reconstitution with exogenous lipids.

Recent developments in detergent-free solubilization methods have emerged as promising alternatives. For instance, styrene-maleic acid (SMA) copolymers can solubilize membrane proteins directly from membranes, forming polymer-wrapped nanoparticles ^12^. However, SMA copolymers are heterogeneous in size, and some membrane proteins may be sensitive to SMA treatment. Additionally, detergent-like lipid-modified peptides have been developed ^13^, such as the chemically modified peptide DeFr_5, which has been shown to solubilize membrane proteins and support cryo-EM studies of the prokaryotic ABC transporter MalFGK2 ^14^. Nevertheless, similar to traditional detergents, these lipid-modified peptides may also replace native lipids due to their lipid components.

Unmodified peptides have also demonstrated the ability to solubilize membrane proteins. For example, NSP and 4F peptides can solubilize phosphatidylcholine (POPC) vesicles ^15^. Detergent-like peptides have been effective in solubilizing olfactory receptors from a cell-free E. coli expression system ^16^. Furthermore, the FFD8 peptide has been utilized to solubilize membranes from Jurkat cells ^17^, while AEM28 and 18A peptides have been shown to solubilize membrane proteins from E. coli membranes ^18^. However, it remains unclear whether unmodified peptides can effectively solubilize eukaryotic membrane proteins on a large scale to facilitate subsequent cryo-EM studies. In this study, we demonstrate that unmodified 4F peptide can solubilize a variety of eukaryotic membrane proteins expressed in FreeStyle 293F cells into peptidiscs. Additionally, we found that the protein solubilized by 4F peptide, exemplified by the human TRPC3 (hTRPC3) channel, can be smoothly converted to MSP-wrapped nanodiscs for high-resolution structural studies.

## Results

### 4F peptide can solubilize tetrameric hTRPC3 channel

To evaluate the effectiveness of peptides in solubilizing eukaryotic membrane proteins, we selected three representative peptides derived from apolipoprotein A-I: 4F ^15^, NSP ^15^, and NSP_r_ ^9^ for solubilization test (Fig. 1A). We also chose the widely used mild detergent LMNG ^19^ as the reference for the solubilization efficiency. For our study, we utilized the homotetrameric human TRPC3 (hTRPC3) channel expressed in FreeStyle 293F cells as a model protein ^20-22^. The hTRPC3 channel is sensitive to harsh detergents, which is evident from the dissociation of the tetrameric peak into a monomeric peak under low-calcium conditions, as observed in fluorescent size-exclusion chromatography ^23^ (Fig. 1B). We found that solubilizing with peptides can effectively solubilize hTRPC3 from the whole-cell membrane, while preserving its tetrameric assembly (Fig. 1B). The solubilization efficiencies of the three peptides were around 20-40 % of that of LMNG (Fig. 1C). Because the solubilization efficiency of these three peptides tested are similar at the same mass concentration and the 4F peptide is much shorter and thus cheaper than the other two (NSP or NSP_r_), we decided to continue our study with 4F peptide due to its cost-effectiveness. We found that the solubilization efficiency of 4F peptide on hTRPC3 channel plateaus at around 2 mg/mL (Fig. 1D). Therefore, we use this concentration for further experiments.

**Figure 1.**
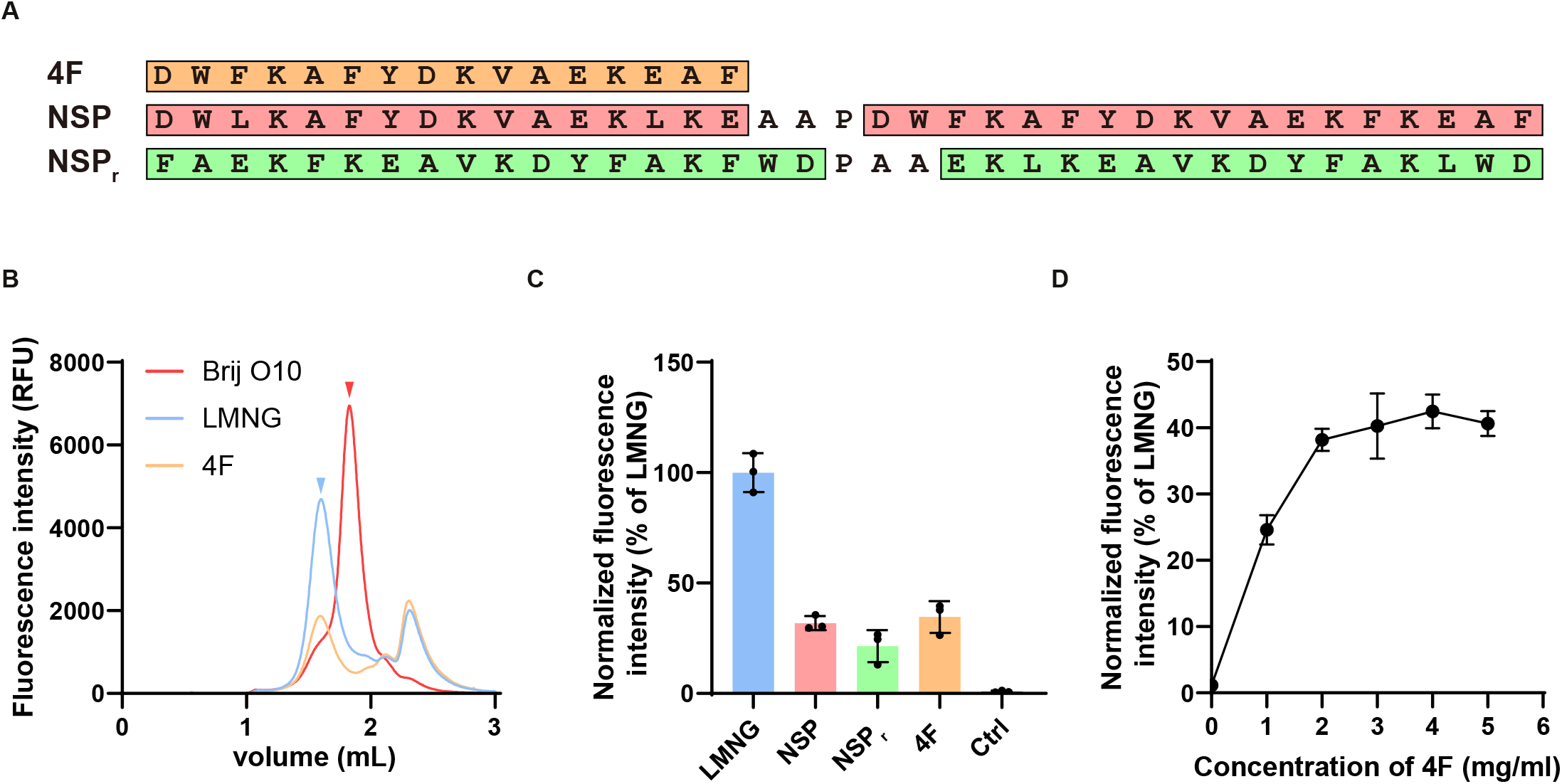
Solubilizing hTRPC3 channel with three representative peptides. A, Sequences of 4F, NSP, and NSP_r_ peptides. The putative alpha-helical sequences are framed and colored. B, Fluorescent size-exclusion chromatogram of the GFP-MBP-tagged hTRPC3 solubilized with Brij O10, LMNG, and 4F peptide under low-calcium conditions. The peak positions for the tetrameric and monomeric hTRPC3 are indicated by blue and red arrows, respectively. C, Relative amount of hTRPC3 solubilized with different peptides. The data are derived from the statistical quantification of the fluorescent signal intensity at the tetrameric hTRPC3 peak position in the fluorescent size-exclusion chromatogram and shown as the mean ± SD. D, The solubilizing efficiency of 4F peptide on hTRPC3 protein. Per mL of cells (∼3×10^6^) is solubilized with 60 µL of 4F peptide at varying concentrations.

### 4F peptide can solubilize various functional eukaryotic membrane proteins

To assess whether the 4F peptide can solubilize other membrane proteins in addition to hTRPC3, we selected a range of eukaryotic membrane proteins, including the human NOX2-p22 complex, the human DUOX1-DUOXA1 complex, human NTCP, and human URAT1, which has been studied in our lab (Fig. 2).

**Figure 2.**
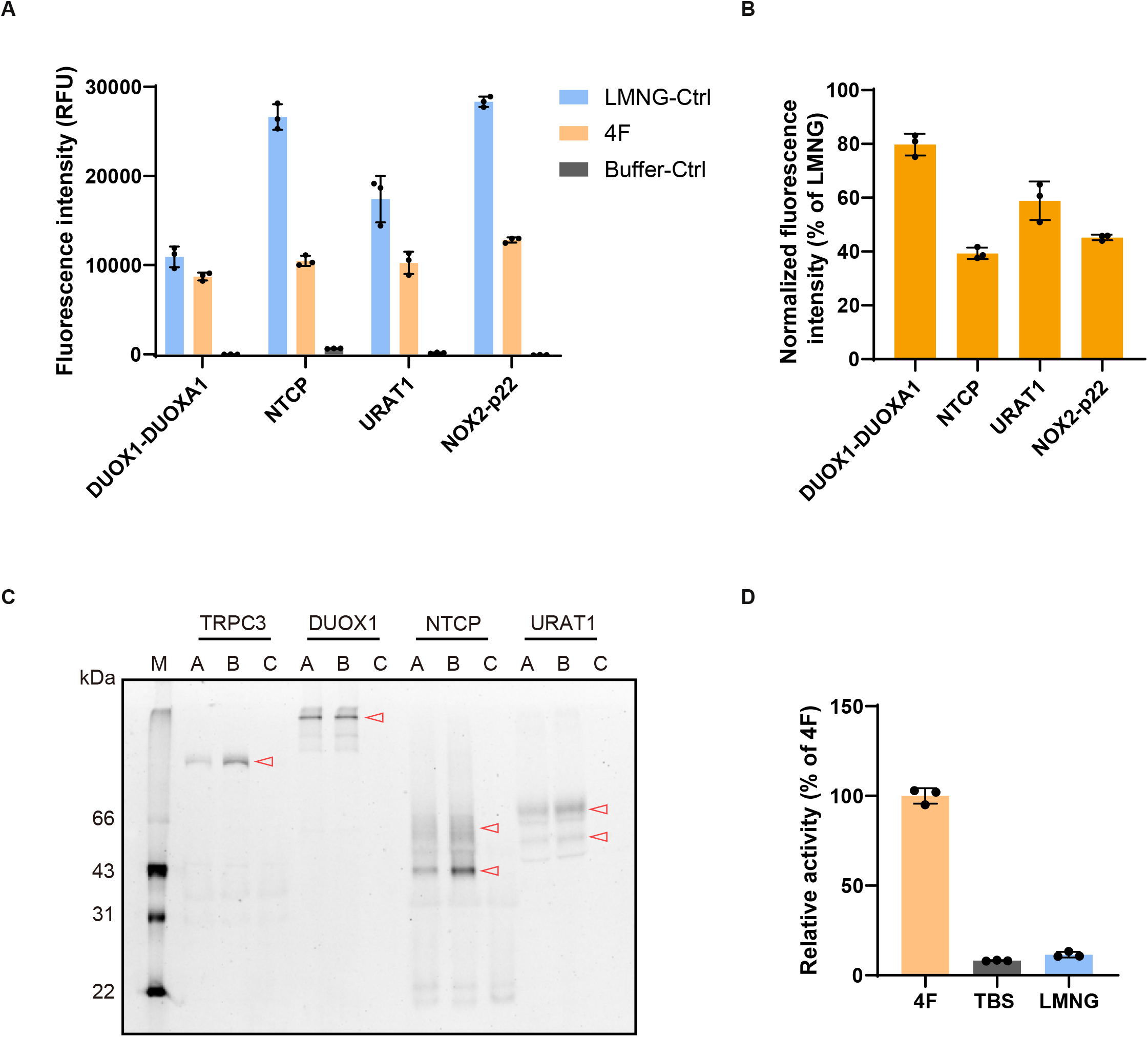
Solubilizing different eukaryotic membrane proteins with 4F peptide. A, Different eukaryotic membrane proteins solubilized with 4F peptide. The data are derived from the statistical quantification of the fluorescent signal intensity at the corresponding protein peaks in the fluorescent size-exclusion chromatogram and shown as the mean ± SD. B, Normalized fluorescent signal intensity of proteins solubilized with 4F peptide (compared with LMNG). C, Fluorescence signals of SDS-PAGE of solubilized proteins (A: 4F; B: LMNG-Ctrl; C: Buffer-control). Red open arrows indicate the positions of target proteins. Some heavily glycosylated proteins (upper bands of NTCP and URAT1) migrate slower than the expected positions. D, The superoxide anion production activity of NOX2-p22 complex solubilized with 4F peptide.

The human NOX2-p22 complex is a heterodimeric complex and serves as the membrane component of the phagocyte NADPH oxidase, which transfers electrons from intracellular NADPH to extracellular oxygen to generate superoxide anions for pathogen elimination ^24^. The activity of the NOX2-p22 complex is sensitive to detergent treatment and highly purified NOX2 in detergent lacks activity ^25,26^. This loss of activity may be attributed to the dissociation of the essential lipidic environment during detergent solubilization, as reconstitution of NOX2-p22 into nanodiscs restores its activity ^27,28^. We found that the 4F peptide can solubilize the NOX2-p22 complex (Fig. 2A-B) while maintaining its activity (Fig. 2D). In contrast, LMNG-solubilized protein has no activity (Fig. 2D). This result showed that solubilization using 4F peptide maintains the lipidic environment that is essential for the activity of NOX2.

The DUOX1-DUOXA1 complex is a tetrameric NADPH oxidase composed of two DUOX1 and two DUOXA1 subunits ^29,30^. We found that 4F peptide can solubilize the DUOX1-DUOXA1 complex, to a similar level compared with LMNG (Fig. 2A-C). Both NTCP ^31-34^ and URAT1 ^35-39^ are SLC transporters, the structures of which have been recently determined. We found that 4F peptide can also effectively solubilize these proteins as well. These results collectively suggest that 4F might be a versatile solubilizing reagent for membrane proteins.

### 4F-peptidisc can be converted to nanodisc

We purified the 4F-peptide solubilized hTRPC3 protein using affinity chromatography with amylose resin, which binds to the maltose-binding protein (MBP) tag on hTRPC3 (Fig. 3). The eluted protein exhibited high purity, as confirmed by SDS-PAGE (Fig. 3B-C), and was further characterized using negative-stain electron microscopy (EM) and dynamic light scattering (DLS) (Fig. 3D). Negative-stain EM revealed the presence of large particles with a diameter of approximately 30 nm (Fig. 3E). DLS analysis indicated a size distribution of particles with a peak at 25 nm, which is larger than hTRPC3 protein itself (Fig. 3D). The significantly larger particle distribution of 4F-solubilized hTRPC3 suggests the co-solubilization and elution of hTRPC3 along with surrounding membrane patches. The highly polydisperse size distribution of the protein eluted from affinity chromatography presents a challenge for structure determination using single-particle cryo-EM, which typically requires particles with a similar size.

**Figure 3.**
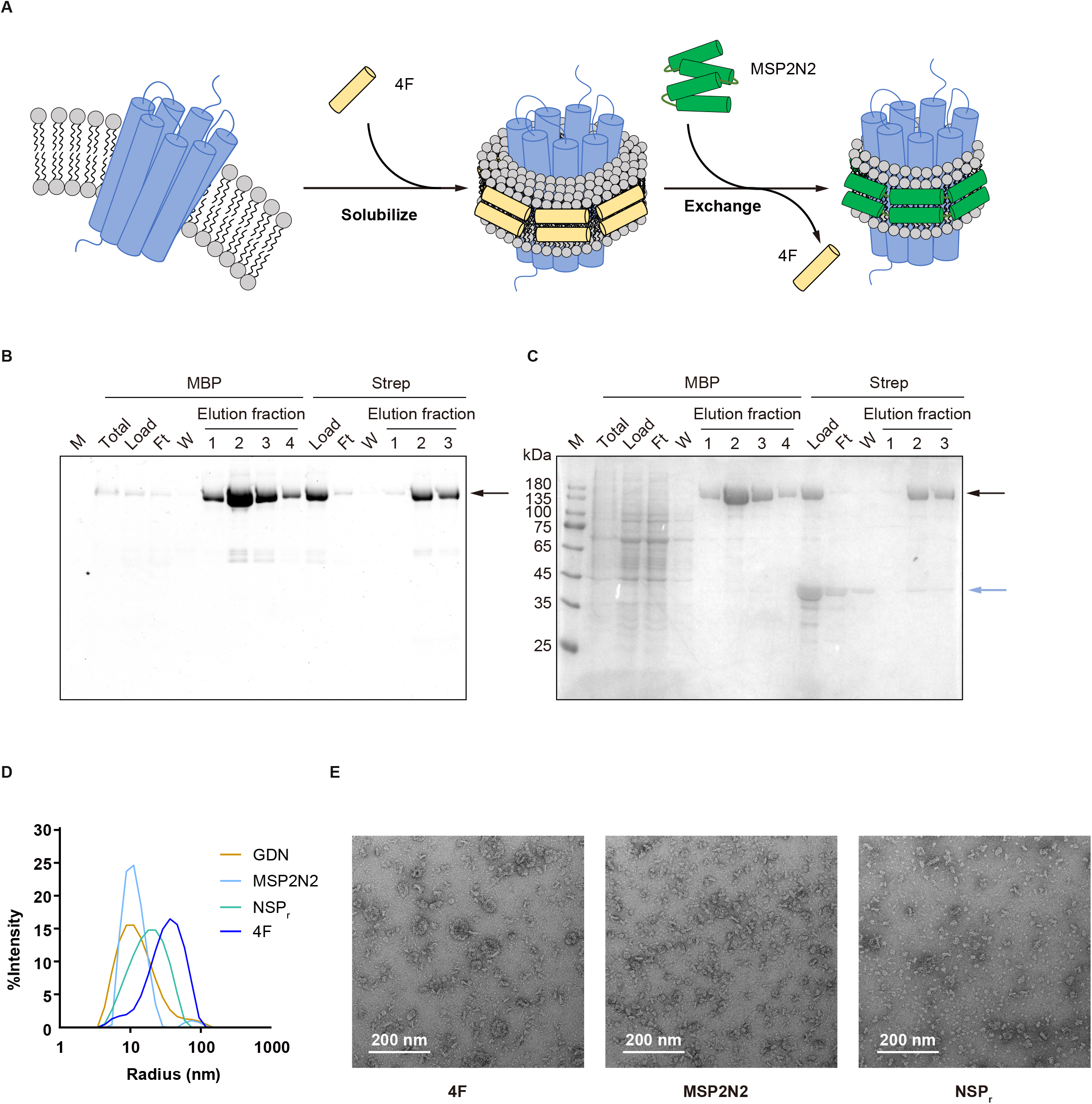
Conversion of hTRPC3 in 4F-peptidisc into nanodisc. A, Model for the conversion process from 4F-peptidiscs to MSP-wrapped nanodiscs. B, Fluorescence SDS-PAGE analysis of the purification processes for MSP2N2-wrapped nanodiscs converted from 4F-solubilized hTRPC3. The black arrow indicates the position of the hTRPC3 bands. C, Coomassie Brilliant Blue staining of the SDS-PAGE gel from (B). The hTRPC3 bands and MSP2N2 bands are indicated by black and blue arrows, respectively. D, The radii of particle determined by DLS. The radii of four purified TRPC3 samples were determined, including the protein in glyco-diosgenin (GDN), the 4F-peptidisc (4F), the MSP2N2-wrapped nanodisc converted from 4F peptidisc (MSP2N2), the NSP_r_ -wrapped peptidiscs converted from 4F peptidiscs (NSP_r_). E, Negative staining images of samples in (D).

To address this issue, we attempted to transfer the hTRPC3 protein from the 4F-peptidisc into MSP-wrapped nanodiscs by incubating hTRPC3 with an excess of MSP2N2. (Fig. 3A) We then further purified the hTRPC3 protein using Streptactin resin, which binds to the Strep tag on hTRPC3, to remove any excessive MSP. The eluted protein from the Strepactin resin contained both hTRPC3 and MSP proteins on SDS-PAGE (Fig. 3B-C), indicating a successful conversion from 4F-wrapped particles to MSP-wrapped nanodiscs. Moreover, negative-stain EM and DLS analyses showed that the MSP-wrapped hTRPC3 nanodiscs exhibited a much smaller size and improved homogeneity compared to the 4F-wrapped hTRPC3 particles, suggesting it might be suitable for cryo-EM studies (Fig. 3D-E).

### Cryo-EM structure of hTRPC3 solubilized using 4F peptide

We supplemented the nanodisc sample with calcium ions and absorbed it onto graphene oxide-coated grids for freezing (Fig. 4A-B). Through single particle analysis, we determined the structure of hTRPC3 at 2.72Å resolution (Fig. 4C-E). We found that the overall structure of hTRPC3 is highly similar to the high-calcium structure determined in detergent previously ^22^. However, closer inspection of the current map reveals a non-protein density surrounded by W322, Y323, K337 on the N-terminal of Pre-S1 elbow, F371, F374 on the S1 helix, and N548, E549 on the loop between S4 and S5 helix, which might represent an endogenous ligand that copurified with hTRPC3 (Fig. 4F). In contrast, a cholesterol hemisuccinate (CHS) density at the same location was observed in the cryo-EM map of previous study, in which hTRPC3 was solubilized with LMNG+CHS detergent ^22^ (Fig. 4G), suggesting that exogenous CHS replaced the endogenous ligand at this site during the membrane solubilization procedure using LMNG+CHS. These results showcase the feasible high-resolution reconstruction of the target membrane protein using 4F-solubilized protein, while preventing the artifact of bound exogenous detergents when using detergent solubilization.

**Figure 4.**
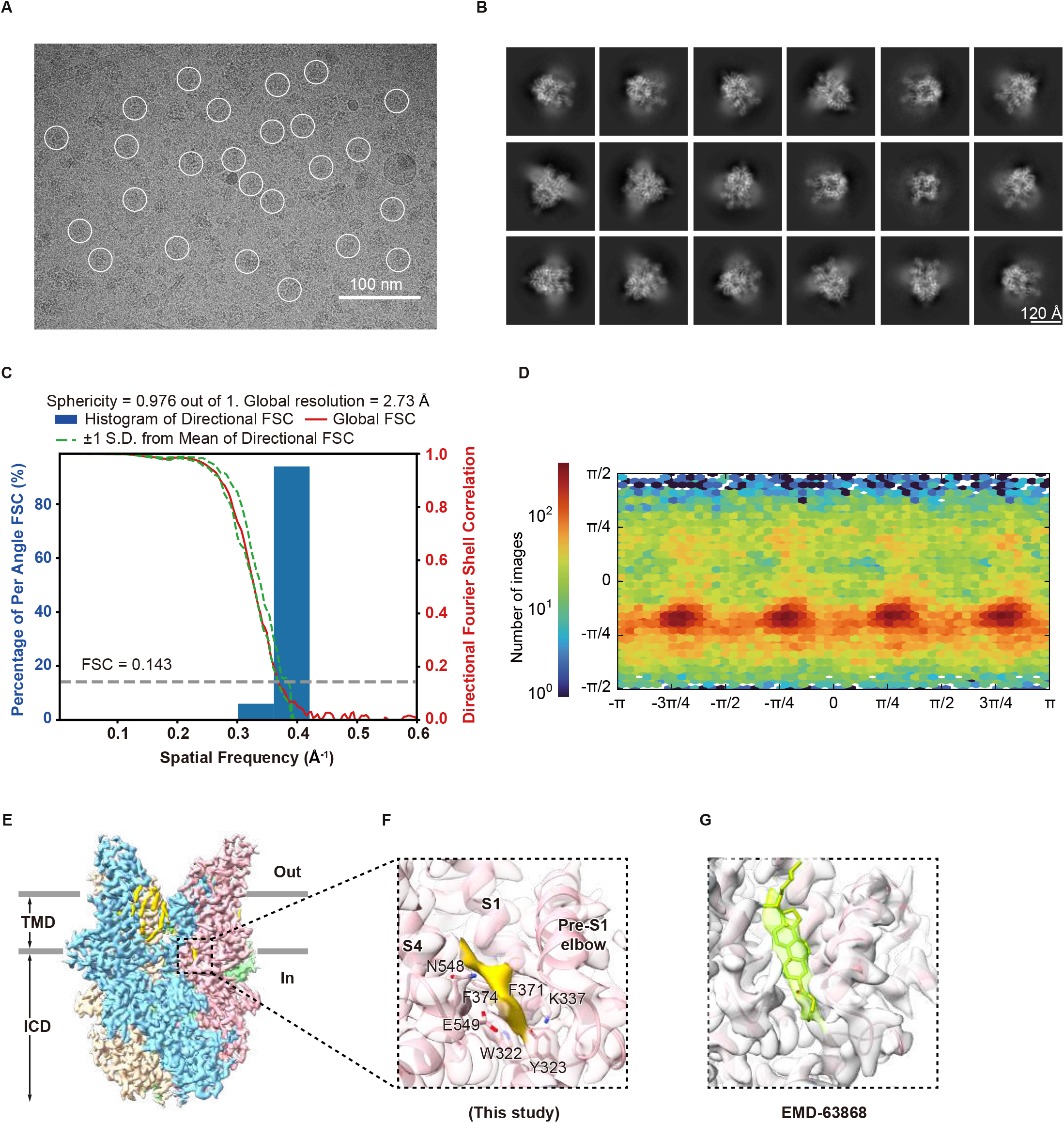
Cryo-EM structure of hTRPC3 solubilized with 4F peptide. A, Representative raw micrograph of hTRPC3 nanodisc converted from 4F-peptidisc. The particles used for final reconstructions are circled in white. B, Representative 2D class average. C, Histogram and Directional FSC Plot for hTRPC3 solubilized with 4F peptide. FSC curves of final refinement for hTRPC3 nanodisc converted from 4F-solublized peptidisc (C4 symmetry) after correction for masking effects. Individual 1D FSC curves are compiled into 3D FSC and represented within a histogram using the 3DFSC server. The spread of the directional resolutions defines by plus and minus one standard deviation from the mean of the directional resolutions. Resolution estimations were based on the criterion of the FSC 0.143 cutoff. D, Angular distribution of final refinement. This is a standard output from cryoSPARC. E, Cryo-EM density map of hTRPC3 nanodisc converted from 4F-solublized peptidisc shown in side view. Four subunits are colored in yellow, blue, pink, and green, respectively. The densities of putative endogenous ligands are colored in gold. F, Densities of the putative endogenous ligand and nearby residues shown in side view. G, Densities of CHS and nearby helixes in hTRPC3 solubilized with LMNG+CHS (EMD-63868). The CHS density is colored in light green.

## Discussion

In summary, our study explored the approach for the solubilization of eukaryotic membrane proteins using the 4F peptide, circumventing the possible pitfalls associated with traditional detergent-based methods. We demonstrated that 4F peptide effectively solubilizes a diverse range of eukaryotic membrane proteins, including the ion channel (hTRPC3), integral membrane enzymes (DUOX1-DUOXA1, and NOX2-p22) and transporters (URAT1 and NTCP), while preserving their structural integrity and activities. The ability to convert 4F-peptidiscs into MSP-wrapped nanodiscs further enhances the structural homogeneity of the particles, enabling high-resolution structural analyses via cryo-EM.

These findings highlight the broad application potential of membrane-solubilizing peptides in membrane protein research. By maintaining the native lipid environment, this approach opens new avenues for understanding the structure and function of integral membrane proteins in their functional states. Future studies could explore the application of this method across a broader spectrum of membrane proteins, thereby advancing our knowledge of their structure and mechanism in native lipid environments.

## Methods

### Materials and Methods

#### Cell Culture

Sf9 insect cells (Thermo Fisher Scientific) were cultured in SIM SF serum-free medium (Sino Biological) at 27 °C. FreeStyle 293F suspension cells (Thermo Fisher Scientific) were cultured in FreeStyle 293 medium (Thermo Fisher Scientific) supplemented with 1% fetal bovine serum (FBS, VisTech), 67 μg/mL penicillin (Macklin), and 139 μg/mL streptomycin (Macklin) at 37 °C with 6% CO_2_ and 70% humidity. The cell lines were routinely checked to be negative for mycoplasma contamination but have not been authenticated.

#### Constructs

The cDNA of hNTCP and hURAT1 were cloned into a modified N-terminal green fluorescent protein (GFP)-tagged pBMCL1 vector as described previously ^40^. The amino acid sequence of hURAT1 was optimized via consensus design method to enhance its expression levels ^39^. Human p22 was fused with a C-terminal GFP tag and cloned into the pBMCL1 vector to generate pBMCL1-p22. hNOX2 was linked into pBMCL1-p22 using LINK method ^41^. N-terminal GFP-tagged hDUOX1 guided by a rat FSHβ signal peptide and C-terminal scarlet-tagged hDUOXA1 were cloned a modified BacMam vector and linked together ^29^. The cDNA of hTRPC3 was inserted into an N-terminal GFP-MBP (Maltose Binding Protein)-tagged pBMCL1 vector for protein purification.

#### Fluorescent size-exclusion chromatography

FreeStyle 293F suspension cells transfected with plasmids for target protein overexpression were harvested by centrifugation at 4000 rpm for 10 minutes at 4°C and washed once with TBS (20 mM Tris pH 8.0 at 4 °C, 150 mM NaCl) buffer. The cell pellet was equally divided into three aliquots, each resuspended in different buffers with equivalent volumes: Buffer A: 5 mg/mL 4F (Genescript,92.5% purity,pH adjusted to 8.0 using Tris pH 8.5 at 4°C prior to usage), 25 mM Tris pH 8.0 at 4 °C and protease inhibitors [1□μg/mL aprotinin, 1□μg/mL pepstatin, 1□μg/mL leupeptin, and 1□mM phenylmethanesulfonyl fluoride (PMSF)] in ddH_2_O; Buffer B: 25 mM Tris pH 8.0 at 4 °C and protease inhibitors in ddH2O; Buffer C: 10mM lauryl maltose-neopentyl glycol (LMNG; Anatrace), 0.1% CHS (Anatrace) and protease inhibitors in TBS buffer.

Following 30-minute incubation on ice, the cell lysates were supplemented with 1 M NaCl to achieve a final concentration of 150 mM (Buffer C group was supplemented with an equivalent volume of TBS buffer instead). Cell lysates were cleared by centrifuge at 40,000 rpm for 30min and supernatants were loaded onto Superose 6 increase 5/150 column (GE Healthcare) for fluorescent size-exclusion chromatography (FSEC) analysis.

#### Enzymatic assay

The superoxide anion-producing activities of the crude cell lysates containing membranes of NOX2-p22 complex were measured using the Amplex Red assay as described before ^27,42,43^. The concentrations of H_2_O_2_ solution were determined by measuring UV-Vis absorbance at 240 nm with a spectrophotometer (Pultton) and calculated using a molar extinction coefficient of 43.6 M^−1^ cm^−1^. The concentration of H_2_O_2_ solution was further validated by reacting with Amplex Red to generate resorufin which has ε_571_ = 69,000 M^−1^ cm^−1^. Then, the H_2_O_2_ solution with known concentration was used to calibrate the standard resorufin fluorescence curve (excitation, 530 nm; emission, 590 nm) measured using a Microplate Reader (Tecan Infinite M Plex) at 30 °C. The reactions were performed at 30°C in a total volume of 90 μl, including 20 μl cleared cell lysates (Prepared as described above), 1 mM MgCl_2_, 1 mM EDTA, 9.3 μM FAD, 500 μM NADPH, 25 μM Amplex Red, 0.067 mg/mL horseradish peroxidase and 0.0576 mg/mL SOD in TBS buffer. 100 nM Trimera was added to activate NOX2 activity. Progress of the reactions was monitored continuously by following the increase of the resorufin fluorescence, and the initial reaction rates were obtained by fitting the curve with a linear equation. The data was processed with Microsoft Excel 2013 and GraphPad Prism 8.0.2.

### Protein expression and purification

The BacMam virus of full-length hTRPC3 was prepared for protein expression as described before ^40^. FreeStyle 293F suspension cells grown in FreeStyle 293 medium at 37 °C with a density of 2.5*10^6^ /mL were infected by BacMam viruses. 10 mM sodium butyrate was added 12 hours post-infection, and the temperature was lowered to 30 °C. Cells were treated with 10 μM U73122 (an inhibitor of phospholipase C) for 10 minutes at 37 °C under standard culture conditions prior to harvesting, then 4 mM EDTA was added to block divalent cations in the culture medium. After 60 hours post-infection, cells were collected and washed once with TBS buffer containing 10 μM U73122, 1mM EDTA and protease inhibitors before being frozen at -80 °C.

For purification of hTRPC3 protein, cell pellets corresponding to 400 mL culture were solubilized in 25 mL lysis buffer (2 mg/mL 4F, 25 mM Tris pH 8.0 at 4 °C, 5 mM DTT, 1 mM EDTA, 10 μM U73122 and protease inhibitors) and rotated at 4 °Cfor 30 min. After centrifugation at 40,000 rpm for 40 min in the Type 50.2 rotor (Beckman), the supernatant was loaded onto a column packed with amylose resin (NEB). The resin was washed with TBS buffer containing 5 mM DTT and 1 mM EDTA. Protein was eluted with MBP elution buffer (80 mM maltose, 100 mM NaCl, 20 mM Tris pH 8.0 at 4 °C, 5 mM DTT, and 1 mM EDTA). After incubation with MSP2N2 at a 1:10 molar ratio for 1 h at room temperature, the collected protein was loaded onto Streptactin resin. Then, the column was washed with TBS containing 5 mM DTT. The target protein was eluted with Strep elution buffer (50□mM Tris pH 8.0 at 4□°C, 100□mM NaCl, 5 mM desthiobiotin, and 5 mM DTT) and collected for cryo-EM studies.

### Dynamic Light Scattering (DLS)

The particle sizes of protein samples were determined using Dynamic Light Scattering (DLS). The DLS experiments were performed on a DynaPro NanoStar instrument (WYATT, CA), operating at λ = 658 nm, produced at a scattering angle of90° at 25 °C. The samples were centrifuged at 40,000 rpm for 30min and supernatants (A_280_ ≈ 0.1) were measured in quartz cuvettes.

### Negative stain electron microscopy

lacey carbon film grids (Zhongjingkeyi Technology Co., Ltd) were glow discharged (31 mA, 30 secs) using PDC-32G-Plasma-Cleaner (Harrick Plasma, Inc). The protein samples (A_280_ ≈ 0.05) were applied onto the grids for 1 minute. After 5 times quickly washed by sterilized ddH_2_O, the samples were stained with 0.75% uranyl formate for 1 minute. Images were collected using a 200kV TEM (JEOL JEM-F200) equipped with a Oneview CMOS camera (Gantan).

### Cryo-EM sample preparation

1 mM CaCl_2_ was added to the purified hTRPC3 nanodiscs with A_280_ ≈ 0.27. After centrifuging at 40,000 rpm for 30 min, the supernatant was collected and supplemented with 0.5 mM fluorinated octyl maltoside (FOM). The protein was added on GIG (1/1) Cu grids coated with graphene oxide ^44^ for cryo-EM sample preparation.

### Cryo-EM data collection

Cryo-EM grids were screened on the Talos Arctica electron microscope (Thermo Fisher Scientific) operating at 200 kV using a K2 camera (Thermo Fisher Scientific). The images of the screened grid were collected on 300 kV Titan Krios (Thermo Fisher) with a K2 Summit direct electron camera (Thermo Fisher Scientific) and an energy filter set to a slit width of 20 eV at a magnification of 60,000× with a pixel size of 0.834 Å and the defocus ranging from -1.5 to -1.8 *μ*m. Super-resolution movies (32 frames per movie) were collected automatically using Serial EM with a dose rate of 22.15 e^-^/pixel/s on the detector and a total dose of 50 e^-^/ Å^2^.

### Cryo-EM data processing

The collected datasets were first gain-corrected, motion-corrected, anisotropic magnification corrected, and dose-weighted by MotionCor2 ^45^. The selected micrographs were then imported into cryoSPARC ^46^. A round of template picking, 2D and 3D classification, and 3D refinement was performed in cryoSPARC. The resulting particles from the best class were used as “Seeds” for seed-facilitated 3D classification of particles. The particles selected were subsequently used for further local CTF and local refinement with C4 symmetry. The resolution estimation was based on the gold standard FSC 0.143 cut-off. Directional FSC was calculated by uploading the map and two half-maps to the 3DFSC server 3dfsc.salk.edu ^47^.

### Model building, refinement, and validation

The hTRPC3 models was manually rebuilt using Coot ^48^ based on the published model 7DXB and further refined using PHENIX ^49^.

### Quantification and statistical analysis

Data processing and statistical analysis were conducted using the software GraphPad Prism. Statistical details could be found in the methods details and figure legends. Global resolution estimations of cryo-EM density maps are based on the 0.143 Fourier Shell Correlation criterion ^50^. The local resolution was estimated using cryoSPARC. The number of independent experiments (N) and the relevant statistical parameters for each experiment (such as mean or standard deviation) are described in the figure legends. No statistical methods were used to pre-determine sample sizes.

## Acknowledgments

Cryo-EM data collection was supported by the Electron microscopy laboratory and the Cryo-EM platform of Peking University with the assistance of Xuemei Li, Zhenxi Guo, Changdong Qin, and Guopeng Wang. Part of the structural computation was also performed on the Computing Platform of the Center for Life Science and High-performance Computing Platform of Peking University. We thank the National Center for Protein Sciences at Peking University in Beijing, China for assistance with negative stain EM. The work is supported by grants from the National Natural Science Foundation of China (32225027 to L.C.) and Center for Life Sciences (CLS to L.C.).

## Author contributions

L.C. initiated the project and wrote the manuscript draft. J. Z., prepared the nanodisc sample and performed the cryo-EM studies. Y.S., W.T., X. L., and W.G. carried out solubilization using 4F peptide. All authors contributed to the manuscript preparation.

## Data availability

Cryo-EM maps and the atomic coordinate of hTRPC3 solubilized with 4F peptide have been deposited in the EMDB and PDB under the ID codes EMDB: EMD-65025 and PDB: 9VFI respectively. Any additional information required to reanalyze the data reported in this work is available from the lead contact upon request.

## Figure Legends

**Figure S1.**
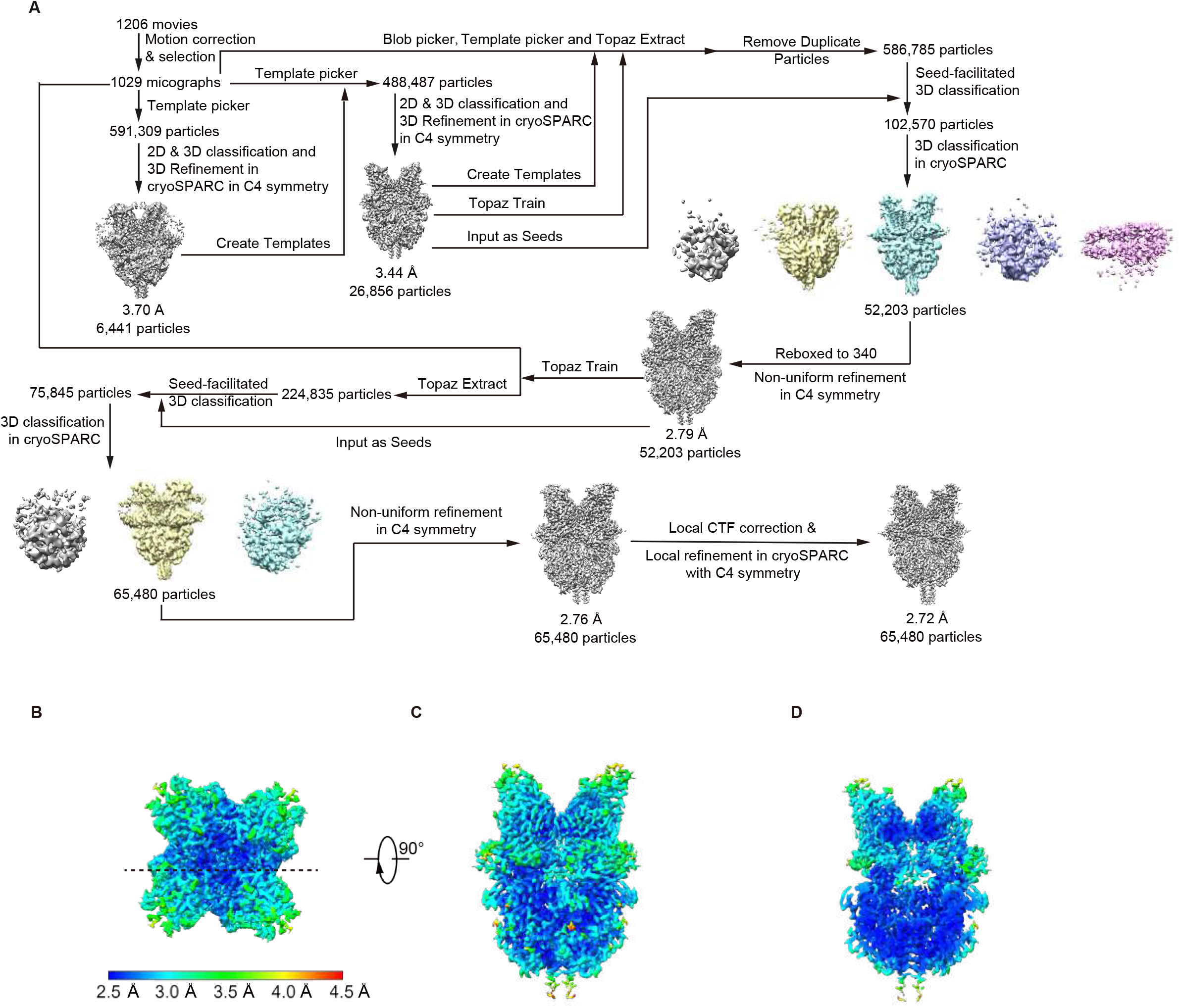
Cryo-EM image analysis of hTRPC3 solubilized with 4F peptide. A, Flowchart of hTRPC3 nanodisc converted from 4F-solublized peptidisc cryo-EM data processing. B, Local resolution estimation of hTRPC3 nanodisc in top view. C, Local resolution estimation of hTRPC3 nanodisc in side view. D, The cross-section of local resolution estimation of hTRPC3 nanodisc. The position of the cross-section was indicated as the dashed line in B.

---

**Table 1.** Cryo-EM data collection, refinement and validation statistics.

|  | PDB ID<br>EMDB ID | hTRPC3 solubilized with 4F peptide<br>9VFI<br>EMD-65025 |
| --- | --- | --- |
| **Data collection and processing** |  |  |
| Magnification |  | 60,000 × |
| Voltage (kV) |  | 300 |
| Electron exposure (e<sup>-</sup>/Å<sup>2</sup>) |  | 50 |
| Defocus range (μm) |  | -1.5 to -1.8 |
| Pixel size (Å) |  | 0.834 |
| Symmetry imposed |  | *C*4 |
| Initial particle images (no.) |  | 512,895 |
| Final particle images (no.) |  | 65,480 |
| Map resolution (Å) |  | 2.72 |
| FSC threshold |  | 0.143 |
| Map resolution range (Å) |  | 250-2.72 |
| **Refinement** |  |  |
| Initial model used (PDB code) |  | 7DXB |
| Model resolution (Å) |  | 2.9 |
| FSC threshold |  | 0.5 |
| Model resolution range (Å) |  | 250-2.9 |
| Map sharpening *B* factor (Å<sup>2</sup>) |  | -103.2 |
| Model composition |  |  |
| Non-hydrogen atoms |  | 23,248 |
| Protein residues |  | 2884 |
| Ligands |  | 16 |
| *B* factors (Å<sup>2</sup>) |  |  |
| Protein |  | 27.43 |
| Ligand |  | 22.66 |
| R.m.s. deviations |  |  |
| Bond lengths (Å) |  | 0.005 |
| Bond angles (°) |  | 0.958 |
| Validation |  |  |
| MolProbity score |  | 1.61 |
| Clashscore |  | 5.37 |
| Poor rotamers (%) |  | 2.64 |
| Ramachandran plot |  |  |
| Favored (%) |  | 98.10 |
| Allowed (%) |  | 1.90 |
| Disallowed (%) |  | 0.00 |

